# Type one protein phosphatases (TOPPs) catalyze EIN2 dephosphorylation to regulate ethylene signaling in Arabidopsis

**DOI:** 10.1101/2025.09.26.678716

**Authors:** Meifei Su, Qianqian Qin, Jing Zhang, Yingdong Li, Ailin Ye, Suoming Wang, Suiwen Hou

## Abstract

TOPPs (Type one protein phosphatases) widely modulate plant hormone signaling and stress responses in plants, but their roles in ethylene signal transduction remain unknown. This study reveals a reciprocal regulatory relationship between TOPPs and EIN2 (Ethylene insensitive 2)-mediated ethylene signaling. We identified that ethylene can induce *TOPPs*’ expression, and *topp1/4/5* triple mutants exhibited partial ethylene insensitivity with reduced EIN3 protein levels and *ERF1* (Ethylene response factor 1) expression. Mechanistically, TOPPs genetically act upstream of EIN2, physically interacting with its carboxyl-terminal domain (EIN2-C) to dephosphorylate the critical S655 residue. This site-specific dephosphorylation promotes EIN2 stability and EIN2-C nuclear accumulation, thereby activating ethylene responses. Notably, transgenic plants expressing phosphorylation-deficient *EIN2*^*S655A*^ displayed constitutive ethylene responses and improved salt tolerance. Further investigation showed that EIN3/EIL1 (EIN3 like 1) transcriptionally activates *TOPPs*’ expression by binding to their promoter, amplifying ethylene signaling accordingly. Together, our finding establishes TOPPs play an important regulatory role in ethylene signal transduction and elucidate a dephosphorylation-switch mechanism in controlling EIN2 function, providing critical insight into the role of EIN2 post-translational modifications in plant stress adaptation.

## Introduction

Type one protein phosphatases (TOPPs/PP1) are major serine/threonine phosphatases in eukaryotes that regulate diverse plant cellular processes^1-3^. Reversible protein phosphorylation is one of the most common PTMs (Post translational modifications) mediated by kinases and phosphatases, precisely controls the stability, activity, and subcellular localization of many core components in the signaling cascade^4-6^. Arabidopsis contains nine TOPP isoforms (TOPP1-TOPP9), with conserved homologs identified in major crops including rice, wheat, and soybean^7-9^. Studies have demonstrated TOPPs serve as a molecular switch for typical phytohormone signaling. TOPP4 destabilizes DELLA proteins via dephosphorylation, modulating gibberellin signaling^10^, and mediates PIN1 (Pin-formed 1) polar localization and endocytic trafficking in pavement cells by regulating PIN1 phosphorylation status in auxin signaling pathway^11^. In wheat, TdPP1 dephosphorylates BES1 (BRI1-EMS-suppressor 1) to mediate BR-regulated root growth^12^. TOPPs also suppress ABA signaling by inhibiting SnRK2 (Sucrose non-fermenting-1-related protein kinase 2) activity^13-14^, while the *Pseudomonas syringae* effector AvrE can interact with TOPPs to remove this inhibition, promoting ABA accumulation and stomatal movement^15^. In addition, the essential functions of TOPPs in plant stress adaptation and immune response are well-established. OsPP1a and TaPP1a mediate salt tolerance in rice and wheat^16,17^, while GmTOPP13 confers drought tolerance in soybean^18^. Under fixed-carbon starvation, TOPPs promote the formation of the ATG1a (Autophagy-related protein 1a)-ATG13a complex by dephosphorylating ATG13a, thereby activating the autophagy pathway^19^. TOPPs are important plant immunomodulators, and the accumulation of SUT1 (Suppressors of *TOPP4-1*, 1) leads to immune activation of *topp4-1* mutant^20,21^. The interaction between TOPPs and KNL1 (Kinetochore scaffold 1) is required for the proper localization of TOPPs to kinetochores to silence the spindle assembly checkpoint in Arabidopsis^22^.

Ethylene is pivotal in plant growth and development, serving as a key signaling molecule for environmental responses^24-28^. Studies elucidating the classical ethylene signaling pathway through Arabidopsis triple-response mutant screens revealed its core regulatory mechanism^29,30^. Without ethylene, CTR1 (Constitutive triple response 1) kinase phosphorylates EIN2 (Ethylene insensitive 2) in ER (Endoplasmic reticulum), stimulating its degradation and consequently hindering ethylene signal transduction^31^. When ethylene is present, CTR1 activity is inhibited, thereby reducing EIN2 phosphorylation and enhancing its stability. The stabilized EIN2 is then cleaved to release its carboxyl-terminal domain (EIN2-C), which functions in cytoplasm and nucleus. Cytoplasmic EIN2-C suppresses translation of E3 ubiquitin ligases EBF1/2 (EIN3 binding F-box protein 1/2) mRNA, thus preventing degradation of the master transcription factor EIN3/EIL1 (EIN3 like 1)^32^. Parallelly, nuclear EIN2-C interacts with ENAP1/2 (EIN2 target protein 1/2) to enhance EIN3/EIL1 transcriptional activity by regulating histone acetylation. Ultimately EIN3/EIL1 activates *ERFs* (Ethylene response factors) expression and ethylene responses^33-35^. Hence, decreasing EIN2 phosphorylation level is essential for ethylene signal relay from ER to nucleus. In addition to the inhibition of CTR1 kinase activity, the reduction of EIN2 phosphorylation should also be attributed to the role of protein phosphatase, but the corresponding phosphatase has not been discovered so far. Our study demonstrates that TOPPs is the first key phosphatase mediating EIN2 dephosphorylation, accordingly regulating ethylene signaling and stress adaptation.

## Results

### TOPPs are involved in ethylene signaling

To investigate the interplay between TOPPs and ethylene signal, we treated WT (Wild-type) plants with the ethylene precursor ACC (1-aminocyclopropane-1-carboxylic acid) and found most *TOPPs* genes were upregulated (fig. S1). GUS staining revealed that ACC treatment significantly induced *TOPPs* promoter activity in hypocotyls, particularly *TOPP1, TOPP4*, and *TOPP5* (Fig. 1A). Next, we analyzed ACC-induced triple responses in *TOPP* mutants (Supplementary information Table 1). All single *topp* mutants (*topp1*-*topp9*) showed no phenotypic difference from WT (fig. S2, A and B). The functional redundancy among TOPPs family members has necessitated the investigation of multiple mutants^19^. Notably, the *topp1/4/5* triple mutant uniquely exhibited partial ethylene insensitivity, whereas other double and triple mutant combinations maintained a WT-like response (Fig. 1, B and C). Conversely, overexpression of *TOPP4* (*TOPP4-OE*) or *TOPP5* (*TOPP5-OE*) lines showed ACC hypersensitivity (Fig. 1, D and E). Furter, native promoter-driven expression of *TOPP1, TOPP4* and *TOPP5* fully rescued the *topp1/4/5* triple mutant phenotype (Fig. 1, F and G). Molecular analysis demonstrated enhanced EIN3 protein accumulation and *ERF1* expression in *TOPP4-OE* lines after ACC treatment, contrasting with significant suppression of these responses in *topp1/4/5* mutants compared to WT (Fig. 1, H and I). These findings establish TOPPs are important regulators of ethylene signaling.

**Fig. 1.**
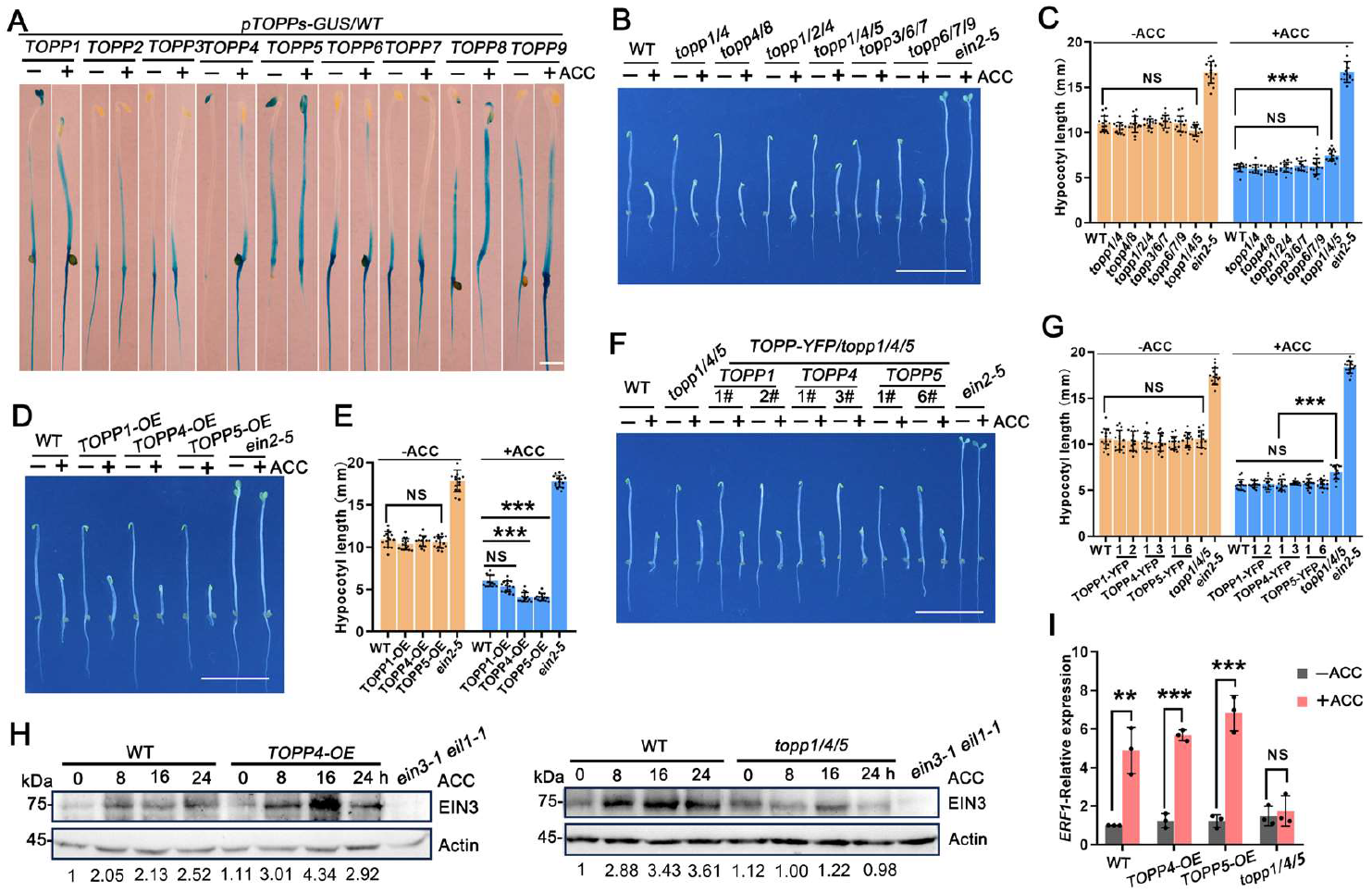
TOPPs are involved in ethylene signaling. **(A**) GUS staining analysis tissue-specific *TOPPs* expression in *pTOPPs-GUS* (*TOPPs* promotor-GUS transgenic plant)*/WT* etiolated seedlings treated with ± 10 μM ACC (8 h). Bar = 100 μm. (**B, D** and **F**) Ethylene-response phenotype of *topp* mutants, *TOPPs-YFP* (*TOPPp:TOPPs-YFP*)*/topp1/4/5* restored lines and *TOPPs-OE* (Overexpressed *TOPPs*) lines treated with ± 10 μM ACC (4 d). Scale bars, 10 mm. (**C, E** and **G**) Hypocotyl measurements of etiolated seedlings. Each bar is the average length ± SD (Standard deviation) of at least 15 hypocotyls per line. (**H**) The EIN3 protein levels were examined in *TOPP4*-*OE* and *topp1/4/5* etiolated seedlings treated with 100 μM ACC for 0, 8, 16 and 24 h using EIN3 antibody, with actin serving as loading control. Relative abundance values shown below blots. (**I**) RT-qPCR analysis of ethylene-responsive gene *ERF1* expression was concurrently performed in *topp1/4/5* and *TOPP4*-*OE, TOPP5*-*OE* etiolated seedlings. Asterisks indicated statistical significance as determined by Student’s *t* test (*\*P*<0.05, *\*\*P*<0.01 and \**\*\*P*<0.001. NS, no significant).

**Fig. 2.**
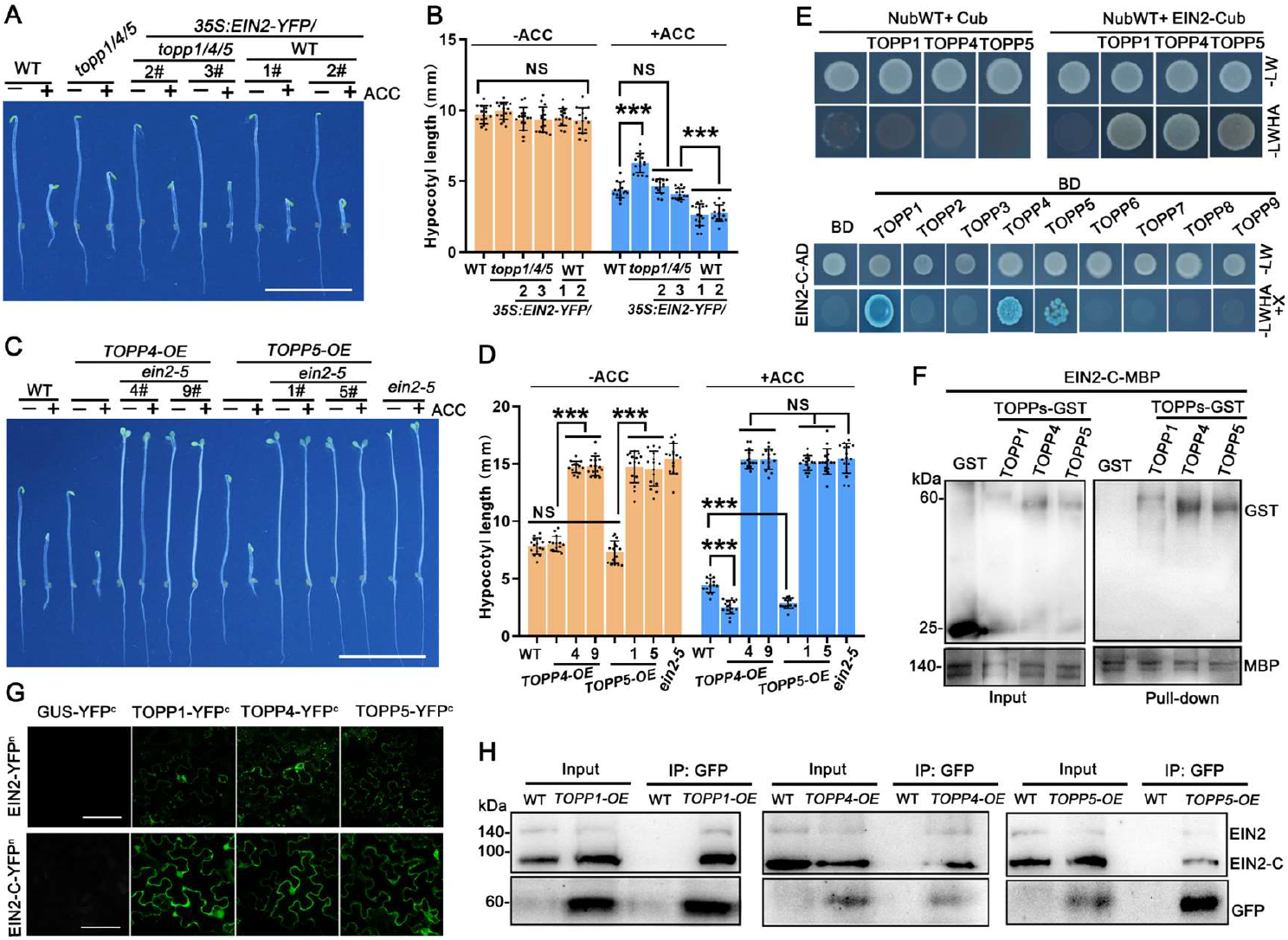
TOPPs are genetically upstream of EIN2 and physically interact with EIN2. (**A** and **C**) Ethylene-response phenotype of the transgenic *35S:EIN2-YFP/topp1/4/5* and *TOPPs-OE/ein2-5* etiolated seedlings treated with ± 10 μM ACC (4 d). Scale bars, 10 mm. (**B** and **D)** Hypocotyl measurements of etiolated seedlings. Each bar is the average length ± SD of at least 15 hypocotyls per line. Asterisks indicated statistical significance as determined by Student’s *t* test (*\*P*<0.05, *\*\*P* <0.01 and \**\*\*P*<0.001. NS, no significant). (**E**) Interaction between TOPPs and full-length EIN2/EIN2-C were analyzed using mbSUS and Y2H assays. Diploid yeast cells were cultured on selective SD media: (-LW) lacking Trp/Leu, (-LWHA) lacking Trp/Leu/His/Ade (with 300 μM Met or X-α-gal). Empty vectors served as negative controls. (**F**) Pull-down assays demonstrating direct binding of TOPPs-GST to EIN2-C-MBP. (**G**) BiFC confirmation of TOPPs-EIN2/EIN2-C interaction in *N. benthamiana*. Scale bars, 50 μM. (**H**) Co-IP validation of TOPPs-EIN2/EIN2-C interaction in Arabidopsis. Total proteins of *TOPPs*-*OE* etiolated seedlings treated with ± 100 μM ACC (16 h) were immunoprecipitated with GFP magnetic beads and analyzed using anti-EIN2. WT was used as a negative control.

### Genetic association and physical interactions between TOPPs and EIN2

To elucidate TOPPs function in ethylene signaling, we performed LC-MS/MS (Liquid chromatograph-tandem mass spectrometry) to analyze the interacting proteins of TOPP4-GFP. We identified two EIN2 peptide segments containing four phosphoserine residues (S645, S648, S655, S659) within the IDR (Intrinsic disordered region) of EIN2-C compared with GFP controls (Table S1 and fig. S3). Notably, TOPP4 emerged as a key EIN2 interactor (data S1), with TOPP1, TOPP4 and TOPP5 co-localized with EIN2 in *N. benthamiana* leaf cells, supporting their functional association (fig. S4A).

Genetic analyses showed that *EIN2* overexpression rescued *topp1/4/5* ethylene insensitive phenotype (Fig. 3, A and B), while the ethylene hypersensitivity phenotype of *TOPP4-OE* and *TOPP5-OE* lines was abolished in *ein2-5* and *ein3-1 eil1-1* backgrounds (Fig. 3, C to D; fig. S, 4B and C). Moreover, the ethylene-triggered EIN3 accumulation was also significantly reduced in *TOPP4-OE/ein2-5* plants (fig. S4D), confirming TOPPs are genetically located upstream of EIN2.

**Fig. 3.**
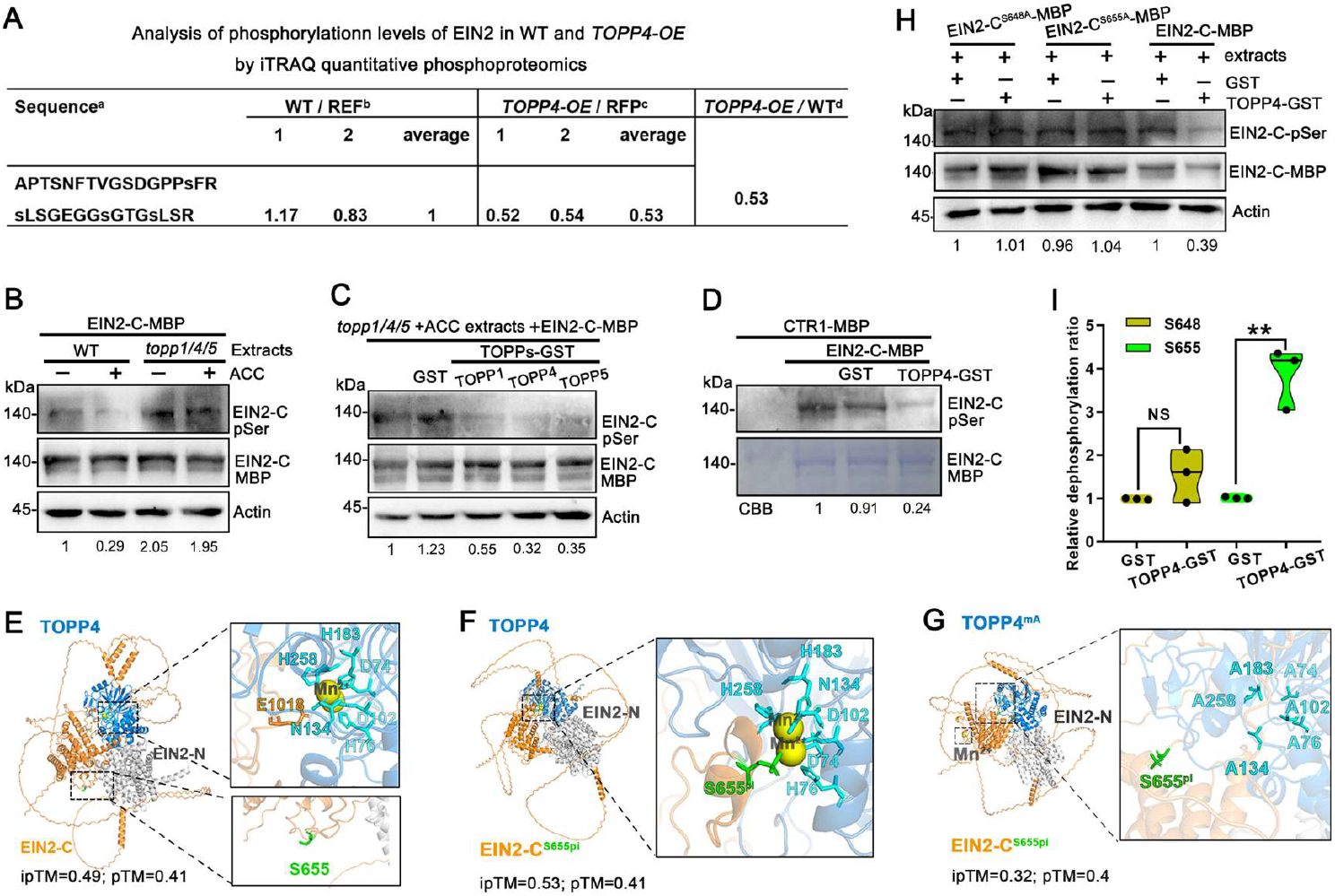
TOPPs specifically dephosphorylate EIN2 at S655 site. (**A**) Peptide abundance ratios (log2-transformed) in WT versus *TOPP4-OE* plants analyzed by iTRAQ quantitative phosphor-proteomics. a, EIN2 phosphopeptide, with phosphorylated residues indicated by lowercase “s”. b– d, Relative abundance of the phosphopeptide: (b) WT vs. internal reference, (c) *TOPP4-OE* vs. internal reference, and (d) *TOPP4-OE* vs. WT. The average value from WT label was used as the internal reference. All ratios represent label-to-reference comparisons. (**B, C** and **H**), *In vitro* phosphorylation/dephosphorylation assays detected EIN2-C-MBP phosphorylation level using anti-pSer immunoblotting (upper panel). Protein loading was confirmed by anti-MBP/CBB staining (middle panel), with GST and actin serving as negative and loading controls (bottom panels). Relative protein levels are shown below each lane. (B) Cell-free kinase assays of equal amounts of recombinant EIN2-C-MBP (as substrate) were incubated with total proteins (as kinase sources) extracted from WT and *topp1/4/5* etiolated seedlings treated with ± 100 μM ACC (16 h), followed by separation with SDS-PAGE. (C and E) Cell-free dephosphorylation assays were performed by supplementing TOPP-GST or GST in kinase assays. (**D**) Dephosphorylation assay of EIN2-C-MBP by TOPPs-GST *in vitro*. (**E** to **G**) The overall structure of EIN2 (EIN2-N, grey. EIN2-C, orange)-TOPP4 (navy blue) complex predicted by AlphaFold 3, TOPP4 catalytic center (D74, H76, D102, N134, H183, H258, cyan). Manganese ions (Mn2+, yellow). (G) Non-phosphorylated EIN2-TOPP4 complex. (H) Phosphorylated EIN2 (S655Pi, green)-TOPP4 complex. (I) Phosphorylated EIN2-TOPP4mA (Mutation of 6 catalytic residues of TOPP4 to A74, A76, A102, A134, A183, A258, cyan) complex. iPTM (Interface prediction template modeling score) and pTM (Predicted template modeling score), showing high interaction confidence (pTM+ipTM >0.75). (**I**) Dephosphorylation of synthesized EIN2 phosphopeptide by GST and TOPP4-GST was analyzed by LC-MS/MS, with S648 and S655 dephosphorylation ratios (dephosphorylated/phosphorylated peptides) determined from three biological replicates. Asterisks indicated statistical significance as determined by Student’s *t* test (*\*P*<0.05, *\*\*P*<0.01 and *\*\*\*P*<0.001. NS, no significant).

We further examined the interactions between TOPPs family proteins and EIN2. *In vitro* analyses using the mbSUS (Mating-based split-ubiquitin system), Y2H (Yeast two-hybrid), and Pull-down experiments revealed direct interactions between TOPP1, TOPP4, or TOPP5 and EIN2, with specific association observed between these TOPP members and EIN2-C (Fig. 3, E to F and fig. S4E). Subsequent *in vivo* validation through BiFC (Bimolecular fluorescence complementation) and Co-IP (Co-immunoprecipitation) assays confirmed these interactions (Fig. 3, G and H). These results demonstrate that TOPPs and EIN2 directly interact to regulate ethylene signaling in Arabidopsis.

### TOPPs dephosphorylate EIN2 at S655 residue

Quantitative phospho-proteomics revealed significantly reduced EIN2 phosphorylation levels in *TOPP4-OE* lines compared to WT (Fig. 3A and data S2). Cell-free kinase and dephosphorylation assays showed ACC-treated *topp1/4/5* mutant extracts significantly enhanced recombinant EIN2-C-MBP phosphorylation levels versus WT (Fig. 3B and fig. S5A), which was reversed by TOPPs-GST supplementation (Fig. 3C and fig. S5B). Additionally, *in vitro* purified CTR1-MBP can phosphorylate EIN2-C-MBP, while TOPP4-GST can dephosphorylate it (Fig. 3D and fig. S5C). Collectively, our data demonstrate that TOPPs serve as a key phosphatase controlling EIN2 dephosphorylation.

We identified four phosphorylation sites (S645, S648, S655, S659) of EIN2 by mass spectrometry (Table S1). AlphaFold 3 predicted distinct EIN2-TOPPs interaction complex (Fig. 3, E and F), Interestingly, in the non-phosphorylated EIN2-TOPP1/TOPP4/TOPP5 complex, S655 lies outside the interaction interface (Fig. 3E and fig. S7, A and B), whereas phosphorylated EIN2 specifically anchors S655 to TOPPs catalytic center (Fig. 3F), comparing to no binding capacity of S645, S648, S659 sites (fig. S7, C to E). Mutating 6 catalytic residues of TOPP4 (TOPP4^mA^) abolished EIN2 S655 binding to its catalytic center (Fig. 3G). Meanwhile, Y2H assays showed that S655 phosphorylation state did not affect their interaction (fig. S7F). These data further suggest that phosphorylated EIN2 facilitates S655 site binding to TOPPs’ catalytic center, enabling its dephosphorylation by TOPPs.

Cell-free dephosphorylation assay showed that alanine substitution at S648 or S655 (EIN2-C^S648A^-MBP or EIN2-C^S655A^-MBP) did not affect EIN2-C-MBP phosphorylation level when incubated with ACC-treated *topp1/4/5* extracts (Fig. 3H and fig. S6A). And addition of TOPP4-GST obviously reduced EIN2-C-MBP phosphorylation, while EIN2-C^S648A^-MBP and EIN2-C^S655A^-MBP phosphorylation levels showed no detectable changes under the same conditions. These results suggest S648 and S655 sites may have important roles in TOPPs regulated EIN2 dephosphorylation. Next we synthesized EIN2 phosphopeptides containing pS648/pS655 (sequence: pSLSGEGGpSGTGSLSR)^36^ and incubated them with TOPP4-GST. Following ultrafiltration removal of the proteins, LC-MS/MS analysis revealed that TOPP4-GST co-incubation specifically increased S655 dephosphorylation radio but had no significant effect on S648 (Fig. 3I and fig. S6B; data 3), confirming that S655 is the key site targeted by TOPPs for EIN2 dephosphorylation.

### Dephosphorylation at S655 site is important for EIN2-mediated ethylene signaling

To assess the impact of TOPPs-mediated dephosphorylation on EIN2 functions, we first performed cell-free degradation assays *in vitro*. Recombinant EIN2-C-MBP stability in ACC-treated *topp1/4/5* mutant extracts was lower than in WT (fig. S8A), while proteasome inhibitor MG132 attenuated this accelerated degradation under the same treatment (fig. S8B). Furthermore, like the effect of alkaline phosphate λpp treatment, TOPP4-GST increased EIN2-YFP and EIN2-C-YFP protein levels in *EIN2-YFP/ein2-5* plants, whereas the Phostop (protein phosphatase inhibitor) reduced them (fig. S, 8C and D), displaying TOPPs enhance EIN2 stability *in vitro*. Meanwhile, *in vivo* analysis detected reduced EIN2 protein levels in *topp1/4/5* mutant versus WT after ACC treatment (Fig. 4A). Consistently, EIN2-YFP and EIN2-C-YFP proteins levels were significantly reduced in *EIN2-YFP/topp1/4/5* plants (Fig. 4B and fig. S8E). It should be particularly pointed out that both EIN2-YFP and EIN2-C-YFP protein accumulated to higher levels in *EIN2*^*S655A*^*-YFP/ein2-5* than in *EIN2*^*S655D*^*-YFP/ein2-5* plants (Fig. 4C). These results indicate that TOPPs stabilize EIN2 through dephosphorylation.

**Fig. 4.**
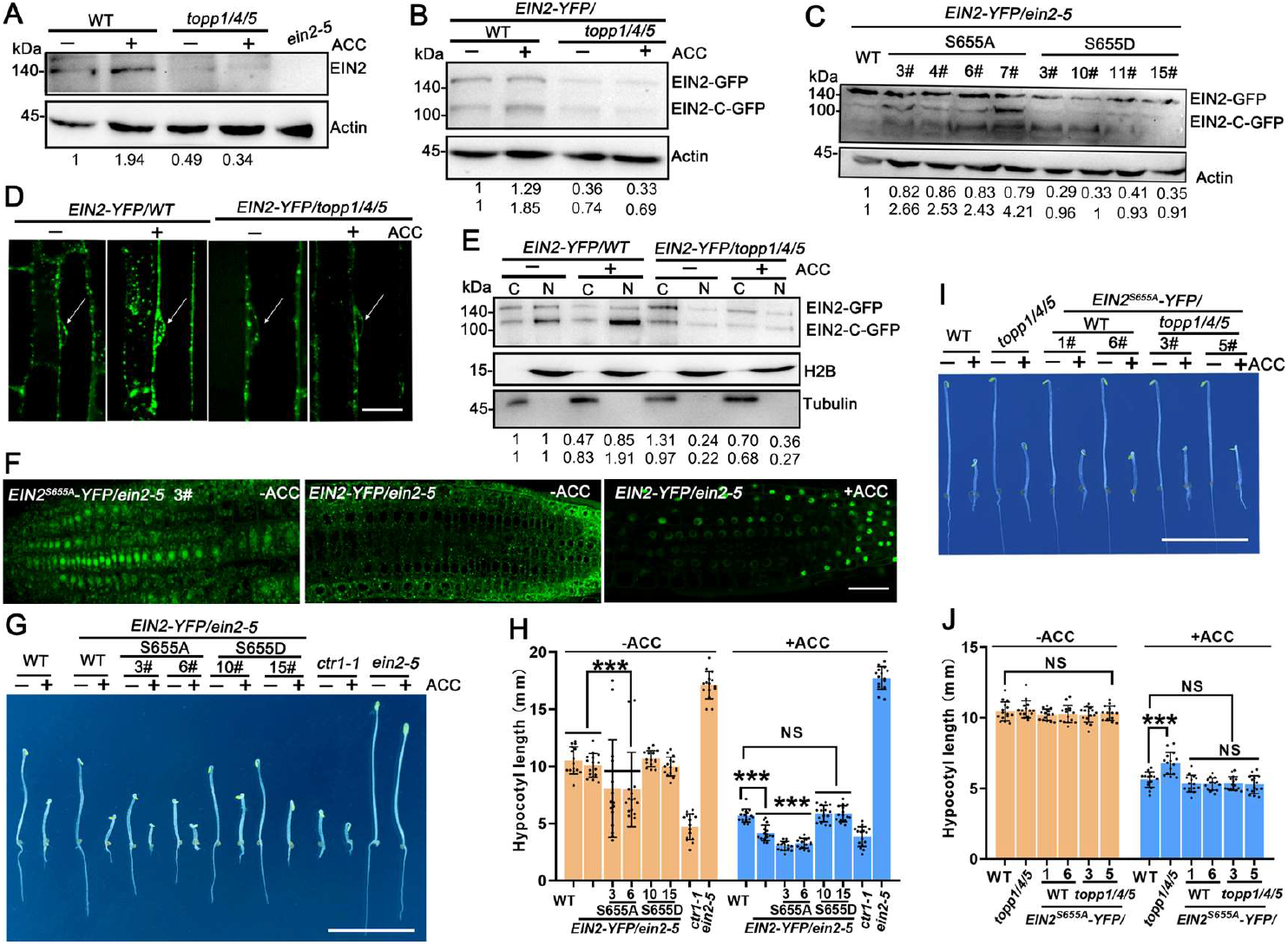
S655 dephosphorylation is important for EIN2-mediated ethylene signaling. (**A**) EIN2 protein levels in WT and *topp1/4/5* mutants treated with ± 100 μM ACC (16 h). Total proteins were subjected to western blotting with anti-EIN2. (**B** and **C**) EIN2-YFP and EIN2-C-YFP levels in *EIN2-YFP/WT, EIN2-YFP/topp1/4/5* and *EIN2*^*S655A*^*-YFP, EIN2*^*S655D*^*-YFP* (*EIN2p:EIN2*^*S655A*^*-YFP* and *EIN2*^*S655D*^*-YFP*) */ein2-5* plants treated with ± 100 μM ACC (16 h), analyzed with anti-GFP, actin served as loading control. (**D**) Nuclear localization of EIN2-YFP (arrows) in roots of *EIN2-YFP/WT* and *EIN2-YFP/topp1/4/5* etiolated seedlings treated with ± 100 μM ACC (16 h). Scale bars, 20 μm. (**E**) Cytoplasmic (C) and nuclear (N) fractions of EIN2-YFP and EIN2-C-YFP in *EIN2-YFP/WT* and *EIN2-YFP/topp1/4/5* seedlings treated with ± 100 μM ACC (16 h). Tubulin and H2B were used as cytoplasm and nucleus loading control, respectively. (**F**) Ethylene-responsive nuclear localization of *EIN2*^*WT*^*-YFP/ein2-5* and constitutive nuclear localization of *EIN2*^*S655A*^*-YFP/ein2-5* in Arabidopsis root cells. Bar = 20 μm. (**G** and **I**) Ethylene-response phenotype of the transgenic *EIN2*^*S655A/D*^*-YFP/ein2-5* and *EIN2*^*S655A*^*-YFP/WT, EIN2*^*S655A*^*-YFP/topp1/4/5* etiolated seedlings treated with ± 10 μM ACC (4 d). Scale bars, 10 mm. (**H** and **J**) Hypocotyl measurements of etiolated seedlings. Each bar is the average length ± SD of at least 15 hypocotyls per line. Asterisks indicated statistical significance as determined by Student’s *t* test (*\*P*<0.05, *\*\*P*<0.01 and \**\*\*P*<0.001. NS, no significant).

ACC treatment could markedly enhance EIN2 fluorescence in nuclear in WT, whereas this phenomenon of increase was not detected in *topp1/4/5* mutant (Fig. 4D and fig. S9A). Nuclear-cytoplasmic fractionation also confirmed ACC-induced nuclear accumulation of EIN2-C-YFP was remarkedly reduced in *topp1/4/5* background compared to WT (Fig. 4E). Notably, *EIN2*^*S655A*^*-YFP/ein2-5* plants displayed constitutive nuclear localization of EIN2-YFP even without ACC treatment, whereas most of EIN2-YFP in *EIN2*^*S655D*^*-YFP/ein2-5* localized in the cytoplasm (Figs. 4F and fig. S9B). In addition, tobacco leaf cells expressing 35S:EIN2^S655A^-C-YFP showed stronger nuclear fluorescence of EIN2-C than EIN2-C^S655D^-YFP (fig. S9C). These findings demonstrate that TOPPs promote EIN2-C nuclear accumulation via S655 site dephosphorylation.

Since phosphorylation state is critical for EIN2 function, we investigated the regulatory role of S648 and S655 site dephosphorylation in ethylene signaling. Transgenic *EIN2*^*S648A*^*-YFP/ein2-5* and *EIN2*^*S648D*^*-YFP/ein2-5* lines showed similar ACC sensitivity (fig. S10, A and B). However, *EIN2*^*S655A*^*-YFP/ein2-5* showed obviously accelerated flowering compared to delayed flowering development in both *EIN2*^*S655D*^*-YFP/ein2-5* and *ein2-5* plants (fig. S11A). Moreover, *EIN2*^*S655A*^*-YFP/ein2-5* showed constitutive ethylene responses under darkness, and ACC treatment also exacerbated its triple-response phenotype (Fig. 4, G and H; fig. S11B), with comparable *EIN2* expression levels across transgenic lines (fig. S11C). In biochemical analysis, EIN3 accumulation and *ERF1* upregulation were observed in the *EIN2*^*S655A*^*-YFP/ein2-5* (fig. S11, D and E). Crucially, expressing *EIN2*^*S655A*^*-YFP* restored ethylene insensitivity phenotype of *topp1/4/5* mutant (Fig. 4, I and J). These results establish that EIN2 dephosphorylation at S655 site is important for activating ethylene signaling.

### EIN2 dephosphorylation is conducive to salt tolerance in Arabidopsis

Ethylene is critical for plant stress adaptation with the ethylene-constitutive mutant *ctr1-1* displays exceptional salt tolerance, and both *ein2-5* and *topp* multiple mutants exhibits similar salt hypersensitivity^14,35^. Consistently, we observed that *TOPP4-OE* plants showed enhanced salt tolerance relative to WT, but this advantage was lost in *ein2-5* and *ein3-1 eil1-1* background with significantly lower fresh weight under salinity stress (Fig. 5, A and B). We evaluated salt stress responses in EIN2 S655 phospho-mutant transgenic lines. Under high-salinity conditions, *EIN2*^*S655A*^*-YFP/ein2-5* plants displayed higher germination rates and cotyledon greening than *EIN2*^*S655D*^*-YFP/ein2-5* (Fig. 5, C to E and fig. S12A). Strikingly, salt-stressed *EIN2*^*S655A*^*-YFP/ein2-5* outperformed other transgenic lines in fresh weight accumulation under 200 mM NaCl treatment (Fig. 5, C and F and fig. S12B). These findings confirm that TOPPs-mediated EIN2 dephosphorylation is conducive to the salt tolerance of plants.

**Fig. 5.**
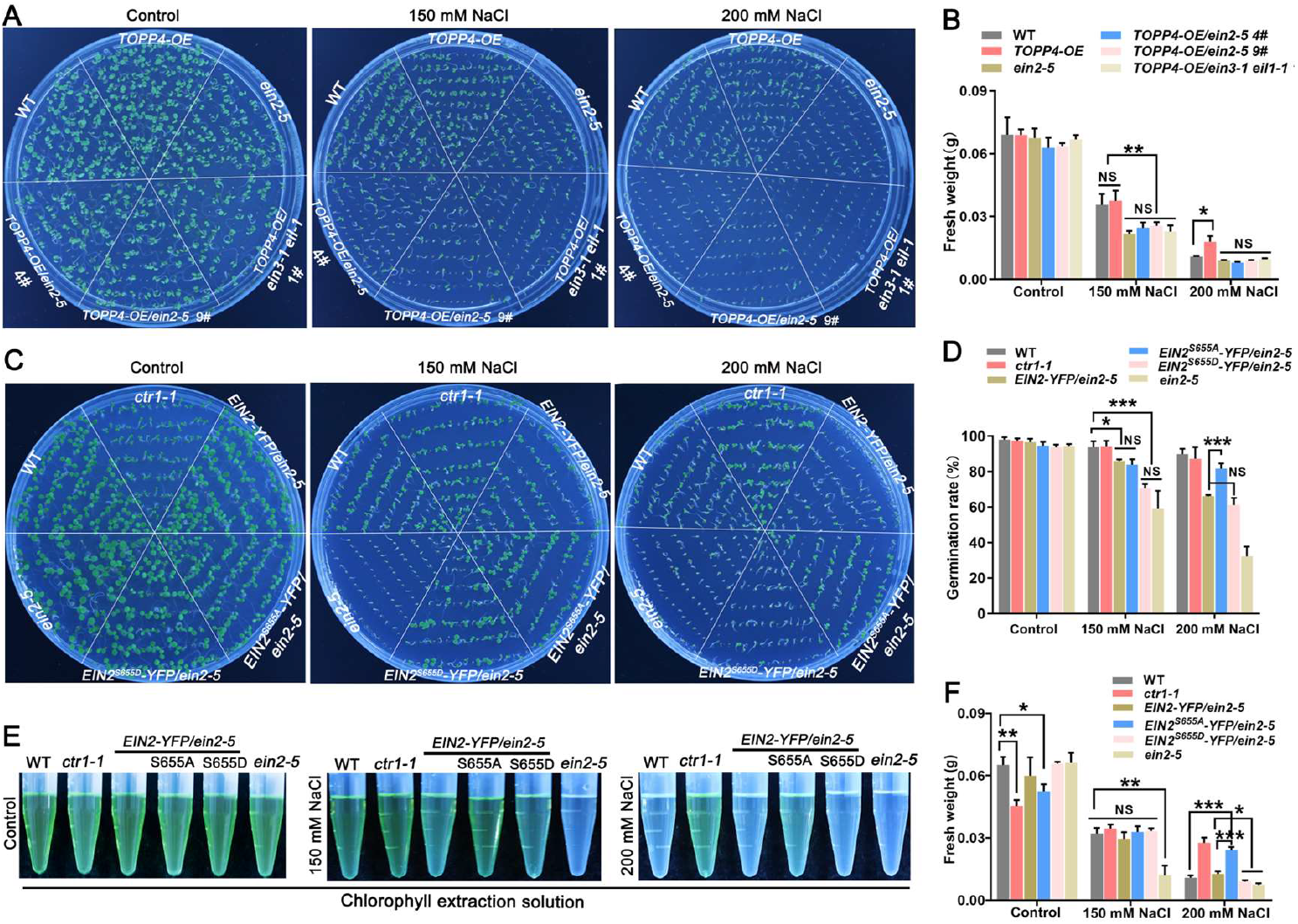
EIN2 dephosphorylation at S655 site is conducive to salt tolerance in Arabidopsis. (**A** and **B**) Salt tolerant phenotype and fresh weight of WT, *TOPP4-OE, ein2-5, TOPP4-OE/ein2-5* and *TOPP4-OE/ein3-1 eil1-1* plants treated with 0, 150, 200 mM NaCl, respectively. (**C** to **F**) Salt tolerant phenotype, seed germination rate, chlorophyll extraction solution and fresh weight of WT, *ein2-5, ctr1-1, EIN2-YFP/ein2-5, EIN2*^*S655A*^*-YFP/ein2-5* and *EIN2*^*S655D*^*-YFP/ein2-5* plants treated with 0, 150, and 200 mM NaCl, respectively. Asterisks indicated statistical significance as determined by Student’s *t* test (*\*P*<0.05, *\*\*P*<0.01 and \**\*\*P*<0.001. NS, no significant).

### Ethylene-induced *TOPPs* expression depends on EIN3/EIL1-mediated transcriptional activation

EIN3/EIL1, the master transcriptional regulator of ethylene signaling, activates target genes through canonical EBS (EIN3-binding site) elements^37,38^. To investigate the mechanism of ethylene-induced *TOPPs* expression, we generated *pTOPPs-GUS/ein3-1 eil1-1* plants by crossing *pTOPPs-GUS/WT* with *ein3-1 eil1-1* mutant. ACC treatment revealed both *TOPP4* and *TOPP5* expressions were significantly inhibited in hypocotyls of *ein3-1 eil1-1* mutant (Fig. 6A and fig. S13A), indicating that EIN3 mediates ethylene-responsive *TOPPs* upregulation. Promoter analysis identified EBS elements TTGTATCTG in *TOPP4* and ATGAAT in *TOPP5* (fig. S13B). Both Y1H (Yeast-one hybrid) and dual luciferase assays confirmed strong interactions between EIN3 and *TOPP4, TOPP5* promoters through these EBS elements (Fig. 6, B and C), which were abolished upon EBS deletion or mutation (Fig. 6B). Further, ChIP-qPCR (Chromatin immunoprecipitation-qPCR) assay using an EIN3 antibody was employed to confirm the binding of EIN3 to the *TOPP4* and *TOPP5* promoters *in planta*. The result showed that EIN3 protein could immunoprecipitate *TOPP4* and *TOPP5* promoter regions containing intact EBS elements in ACC-treated WT plants (Fig. 6D and fig. S13, C and D). Collectively, EIN3/EIL1 can bind *TOPPs* promoter to activate their expression to amplify ethylene signaling in plant.

**Fig. 6.**
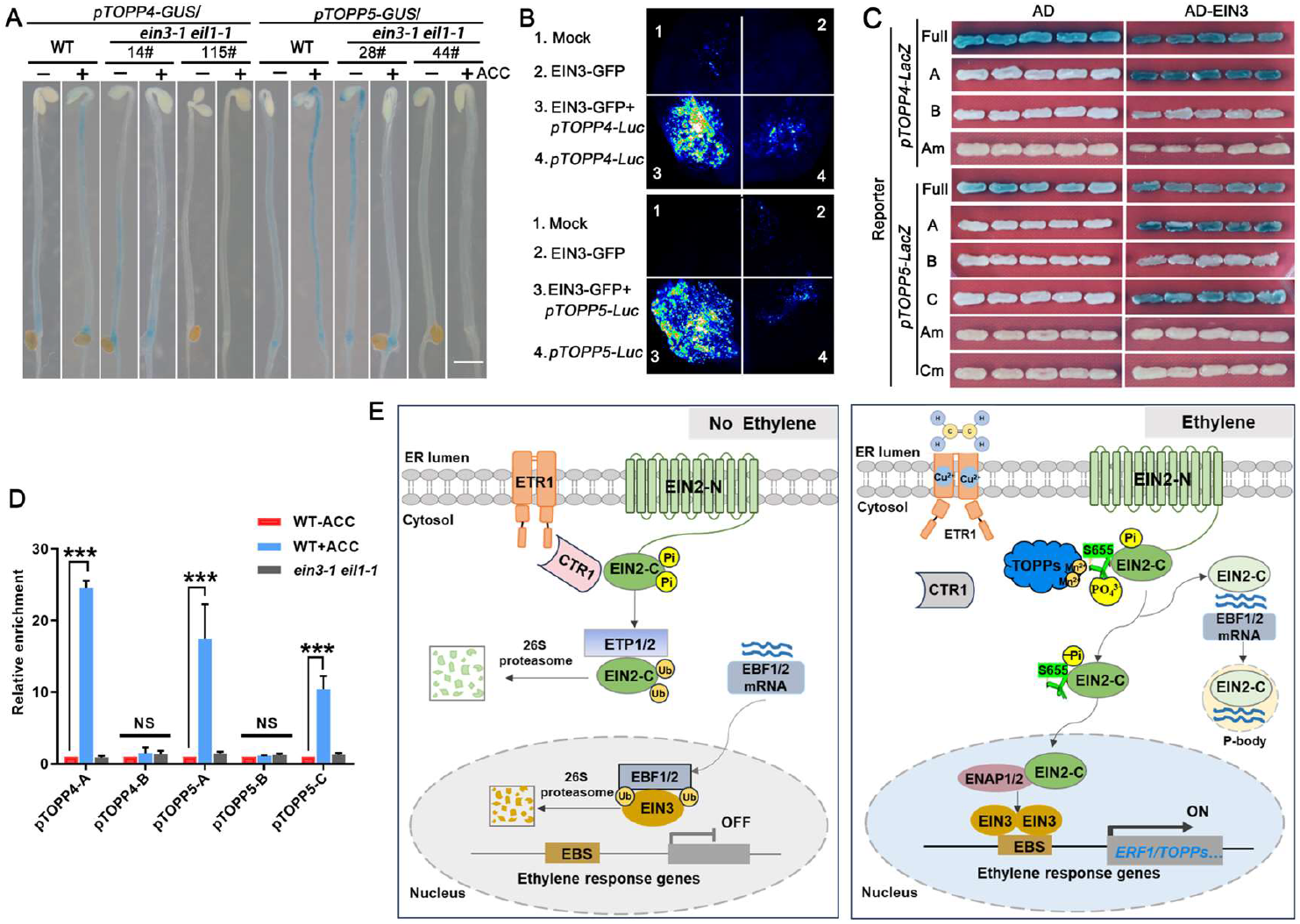
Ethylene induces *TOPPs* expression through EIN3/EIL1-mediated transcriptional activation, establishing a feedback loop with TOPPs-dependent EIN2 dephosphorylation. (**A**) Tissue-specific expression of *pTOPPs-GUS/ein3-1 eil1-1* etiolated seedlings treated with ± 100 μM ACC (8 h). Scale bar, 100 μm. (**B**) Y1H assays demonstrating EIN3 interaction with full-length (full)/truncated (A, B, C)/mutated (Am, Cm) *TOPP4* and *TOPP5* promoters. (**C**) Dual luciferase assays showing EIN3-GFP binding to *TOPP4* and *TOPP5* promoters and enhancing their activity in tobacco leaves. (**D**) ChIP-qPCR assay with EIN3 antibody was performed in WT and *ein3-1 eil1-1* etiolated seedlings treated with ± 100 μM ACC (8 h) to validate EIN3-*TOPPs* promoter interaction.

DNA amounts were normalized to untreated WT. Asterisks indicated statistical significance as determined by Student’s *t* test (*\*P*<0.05, *\*\*P*<0.01 and \**\*\*P*<0.001. NS, no significant). (**E**) working model illustrates how TOPPs regulate ethylene signaling by dephosphorylating EIN2.

When ethylene is absent, receptors-activated CTR1 phosphorylates EIN2, leading to the proteolytic degradation of EIN2 via 26S proteasome and blocking ethylene signaling. When ethylene is present, CTR1 activity is inhibited. Subsequently, TOPPs stabilize EIN2 by binding and dephosphorylating at S655 residue, thereby promoting EIN2-C nuclear accumulation, which enhances EIN3/EIL1 stability to activate *ERF1* and *TOPPs* expression. Ultimately, a self-reinforcing “TOPPs → EIN2→EIN3→*TOPPs*” signal feedback amplification circuit is formed to enhance the ethylene response. Green highlights S655 phosphorylation site. Pi, phosphorylation; PO_4_3-, Phosphate group; -Pi, dephosphorylation; Ub, ubiquitination; Mn^2+^, manganese ions. Arrows indicate signaling progression.

## Discussion

TOPPs serve as central regulatory nodes in plants, orchestrating critical physiological responses through dynamic dephosphorylation of key substrates involved in hormone signaling^10-14^, stress response^16-18^ and immunity^15,20,21^. Ethylene developmental roles are well-characterized^19,20^. Crucially, as the master switch in ethylene signaling, EIN2 activity and subcellular trafficking are highly dependent on phosphorylation status, yet corresponding phosphatases remain unidentified. This study demonstrates that TOPPs directly interact with EIN2 and dephosphorylate it at the S655 site, promoting its stability and EIN2-C nuclear accumulation. This facilitates stable EIN3-mediated transcriptional activation of *ERF1* and *TOPPs*, establishing a self-reinforcing “TOPPs→EIN2→EIN3→*TOPPs*” positive feedback loop that augments ethylene responses (Fig. 6E). Our study uncovers reciprocal interactions between phosphatase TOPPs and ethylene signaling, demonstrating that TOPPs mediate EIN2 dephosphorylation. These findings expand the canonical “receptor-phosphorylation cascade-transcriptional output” framework in ethylene signaling, elucidating precise PTMs regulatory mechanisms in this pathway. Therefore, TOPPs are the first phosphatases found in ethylene signaling.

Consensus holds that each functional TOPPs enzyme consists of a highly conserved catalytic subunit and a regulatory subunit targeting catalytic subunit to specific subcellular compartment, modulate substrate specificity, and trigger specific biological responses^39^. While our study has revealed the critical role of the TOPPs-EIN2 complex in ethylene signaling, the specific regulatory subunits and mechanism controlling TOPPs-mediated dephosphorylation of EIN2 remain to be elucidated. In addition, the *topp1/4/5* mutants exhibit partial ethylene insensitivity compared to the complete ethylene-insensitive mutant *ein2-5*, s indicating that additional factors may contribute to EIN2 dephosphorylation. Arabidopsis possesses around 1000 kinases and 150 phosphatases that form a reversible phosphorylation network^40^. Known regulators of EIN2 phosphorylation include CTR1, TOR (Target of rapamycin), and MtCRA2 (Compact root architecture 2) in plants^34,41,42^, suggesting that other phosphatases are likely involved. Furthermore, since TOPPs-mediated dephosphorylation at S655 site leads to constitutive ethylene signaling, other phosphatases may also target this critical site.

The dynamic stability, subcellular localization, and cleavage of EIN2 into multiple EIN2-C fragments of varying sizes are beneficial to the complexity of its role in ethylene signaling^32,43,44^. PTMs are essential for signaling cascades by precisely regulating central proteins^1^. In Arabidopsis, CTR1-controlled phosphorylation of EIN2 triggers its ubiquitination-dependent degradation^30-34^, with glucose-activated TOR kinase promoting EIN2 phosphorylation to block its nuclear localization^41^, whereas O-glycosylation enhances the stability and nuclear accumulation of SlEIN2 in tomato^45^. Our data reveal that TOPPs-mediated dephosphorylation facilitates EIN2 stability and EIN2-C nuclear accumulation. These PTMs of EIN2 should be intrinsically interrelated, although their precise molecular interplay requires further mechanistic investigation. Importantly, expressing phospho-deficient *EIN2*^*S655A*^*-YFP* in *ein2-5* exhibits enhanced salt tolerance, positioning TOPPs and ethylene as coordinated stress regulators. These findings establish PTMs serve as molecular switches in precise stress-response networks, highlighting their significant potential for stress-resistant crop breeding.

As the proteolytic product of underphosphorylated EIN2, EIN2-C functions in guaranteeing EIN3/EIL1 stability in the ethylene signaling, although it is located both in cytoplasm and nucleus^46^. We found that TOPPs directly dephosphorylate EIN2 at S655 site, facilitating nuclear translocation of most EIN2-YFP, leaving only a small amount of EIN2 in cytoplasm in *EIN2*^*S655A*^*-YFP/ein2-5* lines. How the numerous EIN2-C fragments are sorted into the cytoplasm and nucleus is worthy of further study. In addition, EIN2 phosphorylation sites cluster within the IDR of EIN2-C, a structural feature known to regulate protein structure and function through PTMs^47^. Therefore, exploring the potential relationship among EIN2 dephosphorylation, IDR conformational dynamics and cytoplasmic p-body is of great significance for fully clarifying the spatial regulation of ethylene signaling.

## Materials and Methods

### Plant materials and growth conditions

*A. thaliana* ecotype Columbia was used as the wild-type (WT) control in this study, and all mutants and transgenic plants have been described in <u>Supplementary information Table 2.</u>

### Plant growth conditions and hypocotyl measurement

Arabidopsis and *N. benthamiana* were grown in controlled chambers (22°C, 16/8 h light/dark) under white fluorescent light. For seedling assay, seeds were surface-sterilized and grown vertically on 1/2 MS (Murashige and Skoog) agar plates containing 0 or 10 μM ACC (Sigma-Aldrich). Hypocotyl lengths were quantified from digital images using ImageJ software (NIH).

### RT-qPCR analysis of *TOPPs, EIN2* and *ERF1* mRNA abundance

Four-day-old etiolated seedlings were treated with 10 μM ACC for 8 h. Total RNA was extracted using a Plant RNA Purification Reagent (Omega), followed by genomic DNA removal using DNase I (Promega). Reverse transcription was performed using PrimeScript RT reagent kit (TaKaRa), and quantitative PCR analysis was conducted with SYBR Green Premix Ex Taq II (TaKaRa) on a StepOnePlus Real-Time PCR System. *UBQ10* (Ubiquitin10) served as the endogenous control. The primers used are listed in <u>Supplementary information Table 3.</u>

### GUS staining

Four-day-old etiolated seedlings were treated with or without 10 μM ACC for 8 h, then subjected to GUS staining (50 mM Na-Phosphate, pH 7.0, 1 mM EDTA, 100 mg/mL Chloramphenicol, 2 mM ferricyanide, 1 mg/mL 5-bromo-4-chloro-3-indolyl-b-D glucuroni acid, 0.1% Triton X-100) at 37°C. Following staining, seedlings were destained in 70% ethanol until sufficiently cleared and imaged with a Leica stereomicroscope.

### Protein isolation and immunoblot analysis

For protein extraction, Arabidopsis samples were cryogenically ground in liquid nitrogen and homogenized in protein extraction buffer (50 mM Tris-HCl, pH 7.5, 150 mM NaCl, 5% β-mercaptoethanol, 1 mM EDTA, 10% glycerol, 0.1% NP-40, 1 mM PMSF, 1×Cocktail), after 30 min ice incubation, the lysates were centrifuged at 12,000 ×*g* (4°C, 20 min). The clarified supernatant was immediately mixed with SDS loading buffer at room temperature for subsequent SDS-PAGE analysis.

### Mass spectrometry assay

To prepare samples for mass spectrometry analysis, at least 2 g transgenic material seedlings were collected. Target protein complex was immunoprecipitated with anti-GFP beads (D153-11, MBL), and the immunoprecipitates were separated by SDS– PAGE. Subsequently, the sample requires a proteolytic step with trypsin before injecting to a mass spectrometer. Samples were analyzed using an Orbitrap Fusion Lumos mass spectrometer (Thermo Fisher Scientific) connected to an EASY-nano-LC system. Raw files were analyzed together using Maxquant (1.6.2.10).

### Mating-based split-ubiquitin system and Y2H assay

TOPPs coding sequences were cloned into pX-NubWTgate vector (THY.AP5 strain), while full-length EIN2 coding sequence was inserted into pMetYCgate vector (THY.AP4 strain), with empty vectors as negative controls. Positive transformants were selected on SD/-Trp-Leu medium and protein interactions assayed on SD/-Trp-Leu-His-Ade medium. Yeast colony growth was examined following 4 d incubation at 30°C. For Y2H assay, the carboxyl-terminal domain of EIN2 was fused to the AD vector, while cDNAs of TOPP superfamily genes were cloned into the BD vector. Both constructs were co-transformed into yeast strain Y2H Gold. Transformants were selected on SD/-Trp-Leu medium at 30°C for 48 h. Protein-protein interactions were assessed by growth on SD/-Trp-Leu-His-Ade plates supplemented with X-α-gal (20 mg/mL) under identical conditions.

### *In vitro* protein expression and pull-down assay

The target proteins were expressed in *Escherichia coli* BL21 cells using an IPTG-inducible vector. Post-induction, cells were lysed via sonication in binding buffers (GST: 140 mM NaCl, 2.7 mM KCl, 10 mM Na_2_HPO_4_, 1.8 mM KH_2_PO_4_, pH 7.3; MBP: 20 mM Tris-HCl, 200 mM NaCl, 1 mM EDTA, pH 7.4), and clarified by centrifugation 12,000 × *g* (30 min, 4 ° C). Proteins were affinity-purified using either Glutathione Sepharose 4B beads (GE Healthcare) for GST-tagged proteins or PurKine™ MBP-Tag Dextrin Resin 6FF (Abbkine) for MBP-tagged proteins. After washing with tag-specific buffers (GST: 50 mM Tris-HCl, 10 mM glutathione, pH 8.0; MBP: 20 mM Tris-HCl, 1 mM EDTA, 10 mM maltose, pH 7.4).

Purified bait protein (GST-tagged) was immobilized on glutathione-sepharose beads by incubating at 4°C for 1 h. After blocking with BSA, prey protein (MBP-tagged) was added and incubated for 2 h in binding buffer (20 mM HEPES pH 7.4, 150 mM NaCl, 0.1% Triton X-100). Beads were washed three times to remove unbound proteins, then boiled in SDS-loading buffer. Eluted complexes were analyzed by SDS-PAGE and Western blotting using tag-specific antibodies (anti-GST/anti-MBP) to confirm interaction. The antibodies used are listed in <u>Supplementary information Table 4.</u>

### Bimolecular fluorescence complementation (BiFC) and fluorescence microscopy

For BiFC-based interaction mapping, truncated or full-length EIN2 and TOPPs were fused with nYFP and cYFP, respectively. Constructs were transformed into *Agrobacterium tumefaciens* strain GV3101 and co-expressed in *Nicotiana benthamiana*. For confocal imaging, leaf sections (2×2 mm) were harvested 36-48 h post-infiltration and examined using a Nikon A1+ confocal laser scanning microscope with 20 × objectives. For live-cell imaging, root meristematic cells from four days old Arabidopsis seedlings were observed using a Nikon confocal microscope with a 40× water objectives. YFP was excited with a 488 nm laser and detected between 500 and 550 nm.

### Co-immunoprecipitation (Co-IP) assay

The Co-IP assay was performed as previously described^19^. In brief, four-day-old etiolated seedlings (WT or *TOPPs-OE* transgenic) were lysed in ice-cold IP buffer (10 mM HEPES pH 7.5, 100 mM NaCl, 1 mM EDTA, 10% glycerol, 0.5% Triton X-100, 1× protease cocktail). Lysates were incubated with anti-GFP magnetic beads at 4°C for 3 h, followed by three cold IP buffer washes. Proteins were separated by SDS-PAGE and analyzed by immunoblotting.

### *In vitro* dephosphorylation and cell-free phosphorylation/dephosphorylation assay

CTR1-MBP was incubated with recombinant EIN2-C-MBP in kinase assay buffer at 30°C for 1 h, followed by reaction termination with 10 mM EDTA. Purified TOPP-GST or GST was then added to the mixture and incubated for 30 min. Reactions were halted with 1× SDS-PAGE loading buffer. Phosphorylated EIN2-C-MBP was detected via immunoblotting using an anti-pSer antibody (Abcam, ab9332; 1:1000 dilution). Protein bands were visualized by CBB staining.

For cell-free phosphorylation assays, four-day-old etiolated seedlings of WT and *topp1/4/5* mutants treated with or without ACC, were flash-frozen in liquid nitrogen. Total proteins were extracted using kinase buffer (25 mM Tris-HCl pH 7.5, 50 mM NaCl, 1 mM DTT, 0.1% Triton X-100, 5 mM MgCl_2_, 5 mM MnCl_2_, 1 mM PMSF, 1× protease inhibitor cocktail). EIN2-C-MBP was incubated with seedling extracts in kinase buffer at 30°C for 45 min, followed by reaction termination with 10 mM EDTA. For dephosphorylation, recombinant TOPP-GST or GST (control) proteins were added to phosphorylated EIN2-C-MBP (or its C^S648A/CS655A^ variants) in phosphate buffer (50 mM HEPES pH 7.4, 0.1% Triton X-100, 1 mM NaCl, 0.1% DTT, 5 mM EDTA, 1 mM ATP, 10 mM MnCl_2_) and incubated at 30°C for 90 min. Reactions were halted by adding SDS loading buffer and boiling for 5 min. Proteins were resolved on 10% SDS-PAGE gels and immunoblotted using anti-pSer antibody.

### AlphaFold 3 analysis

Protein interaction between EIN2 and TOPP1, TOPP4 and TOPP5 was predicted by AlphaFold 3 model (https://alphafoldserver.com/). Full protein sequences of EIN2, TOPPs and 2x Mn^2+^ were entered as input for the prediction. Final models were imported and modified with PyMOL V3.1.0 (https://www.pymol.org/). iPTM (Interface prediction template modeling score) and pTM (Predicted template modeling score), showing high interaction confidence (pTM+ipTM >0.75).

### Cell-free degradation assay

Total proteins were extracted from four-day-old WT and *topp1/4/5* etiolated seedlings treated with or without ACC were frozen in liquid nitrogen, and total proteins were extracted with 1×degradation buffer (50 mM Tris-HCl pH 7.5, 150 mM NaCl, 10 mM MgCl2, 1 mM EDTA, 10 mM NaF, 2 mM Na3VO4, 25 mM β-glycerophosphate, 10% (w/v) glycerin, 2 mM DTT, 1 mM PMSF and 1×Cocktail). Each individual assay used recombinant EIN2-C-MBP protein and 1 mM ATP incubated in 100 μL total proteins extracts, then incubated at 28°C and sampled at the indicated intervals for western blotting.

### Dual luciferase assay

Dual-luciferase assays were performed to analyze protein-DNA interactions. Full-length EIN3 was fused to eGFP, while the *TOPPs* promoter sequence were cloned upstream of the Luc (Luciferase) reporter gene. These constructs were co-transformed into *Nicotiana benthamiana* leaves for transient expression analysis.

### Yeast one-hybrid assay

For the yeast one-hybrid assay, AD-fusion constructs and LacZ reporters were co-transformed into the EGY48 yeast strain. Transformants were selected and cultured on SD/-Trp-Ura medium. Yeast transformation and liquid assays were performed as described in the Yeast Protocols Handbook (BD Clontech).

### ChIP-qPCR

Chromatin isolation was performed using four-day-old WT and *ein3-1 eil1-1* etiolated seedlings treated with or without ACC for 8 h. After resuspension, chromatin was sonicated at 4°C to produce 200-600 bp fragments. Immunoprecipitation was performed using monoclonal anti-EIN3 antibody, followed by washing, reverse crosslinking, and DNA amplification. About 10% of sonicated but non-immunoprecipitated chromatin was reverse cross linked and used as an input DNA control. Both immunoprecipitated DNA and input DNA were analyzed by RT-qPCR (Applied Biosystems).

## Acknowledgments

We thank Prof. Xiaoping Gou (Lanzhou University) providing mbSUS related vectors. Prof. Fang Lin (Lanzhou University) providing Y1H and dual luciferase assay related vectors. Prof. QiJun Chen (China Agricultural University) for pCBC-DT1T2 and pHEE401 vector. We are grateful to Liang Peng, LiPing Guan, and YaHu Gao (Core Facility at Life Science Research, Lanzhou University) for technical assistance.

## Data and materials availability

All data needed to evaluate the conclusions in the paper are present in the paper and/or the Supplementary

